# New insights into sperm ultrastructure through enhanced scanning electron microscopy

**DOI:** 10.1101/2020.11.12.380808

**Authors:** Denis Korneev, D. Jo Merriner, Gediminas Gervinskas, Alex de Marco, Moira K O’Bryan

## Abstract

The analysis of spermatozoa morphology is fundamental to understand male fertility and the aetiology of infertility. Traditionally scanning electron microscopy (SEM) has been used to define surface topology. Recently, however, it has become a critical tool for three-dimensionally analyse of internal cellular ultrastructure. Modern SEM provides nanometer-scale resolution, but the meaningfulness of such information is proportional to the quality of the sample preservation. In this study, we demonstrate that sperm quickly and robustly adhere to gold-coated surfaces. Leveraging this property, we developed three step-by-step protocols fulfilling different needs for sperm imaging: chemically fixed monolayers for SEM examination of the external morphology, and two high-pressure freezing-based protocols for fast SEM examination of full cell internal morphology and focused ion-beam SEM (FIB-SEM) tomography. These analyses allow previously unappreciated insights into mouse sperm ultrastructure, including the identification of novel structures within the fibrous sheath and domain-specific interactions between the plasma membrane and exosome-like structures.

## Introduction

Sperm ultrastructure and the relationship between sperm structure and function is a primary determinant of male fertility, and thus infertility, and a critical driver in evolutionary processes (1). Accurately describing sperm morphology requires excellent visualisation tools. Spermatozoa were first visualised with electron microscopy (transmission) in the 1940s (2, 3). Scanning Electron Microscopes (SEM) became commercially available in the late 1960s, and the first studies of sperm external morphology using SEM appeared around the same time. Since this time, there have been significant advances in SEM technologies (4). The analysis of male gametes has not, however, keep pace with these technologies, thus creating a significant knowledge gap. The ability to fill this gap has the potential to inform gene function, the aetiology of male infertility, sperm behaviour within the female reproductive tract and how subtle changes in sperm structure-function may drive evolutionary processes. Towards this goal, we developed a fast and reliable protocol of sperm monolayer preparation, which can be used in chemical fixation (SEM examination of external morphology) as well as in cryogenic preservation method (high-pressure freezing and freeze substitution) allowing the examination of internal morphology at electron microscopic resolution.

SEM imaging is more efficient and provides better results if the cells are distributed as a monolayer. This is especially true for highly elongated cells such as spermatozoa which easily overlap during the preparation. The most straightforward procedure to prepare a cell monolayer from a suspension consists of placing a droplet on a surface with high adhesive properties, allowing the cells to settle, and then remove the residual liquid. Here, the adhesion of the cells to the surface must be strong enough to prevent their detachment during the processing steps. Sperm preparation protocols developed in the early 1970s (5–7) consisted of the sequential use of chemical fixatives (e.g. formaldehyde, glutaraldehyde and osmium tetroxide), washing steps with saline solutions, the formation of a cell monolayer, dehydration (either by air, critical-point carbon dioxide, or chemical), and finally coating with an electrically conductive layer such as metal or carbon. At the time, poly-lysine coating of plates and grids became the preferred method for cell monolayer preparation for electron microscopy examination (8). Although general morphology can be examined using the described approach, the ultrastructural preservation is extremely poor. Such poor ultrastructural preservation has become increasingly obvious, and problematic, over the past two decades, as a consequence of the significant improvements in SEM imaging, which is now routinely capable of nanometre resolution of the cell topography (9). Given the tight relationship between the sperm structure and functionality, it is crucial that sample preparation and analysis protocols are revised (10).

Focused ion beam scanning electron microscopy tomography (FIB-SEM) uses a flux of accelerated ions to serially remove thin layers (down to 2 nm (11)) from the face surface of a sample (in our case a resin block) and sequentially image the newly exposed surface using the SEM. This technique has been intensively developed in the last two decades and fills the “gap” between TEM tomography and light microscopy. FIB-SEM tomography makes it possible to visualise relatively large (50 × 50 × 50 μm and even larger) biological structures with nanoscale isotropic volume resolution (typically ranging from 4 to 20 nm) (12). This innovative technique provides a unique possibility to examine sperm morphology using a full-cell 3D-reconstruction. A proper design of sample preparation is crucial to introduce FIB-SEM tomography in reproductive biology studies.

In this manuscript, we describe a step-by-step protocol to prepare sperm monolayers for high-resolution SEM imaging of the external and internal cellular morphology. We further describe a step-by-step workflow to prepare and embed sperm monolayers for fast FIB-SEM tomography. Collectively, these protocols allow the visualisation of previously unappreciated structures within and on sperm.

## Results

### 1. External sperm morphology

Within this study, we aimed to develop a protocol that would minimise sample handling, and thus maximise structural integrity, and provide a flexible and technically accessible protocol for use by sperm biologists. We also aimed to reassess what is known about mouse sperm morphology. This required returning to first principals for several aspects of the method. An ideal sample for high-resolution SEM imaging should be dry, conductive, stable under the electron beam (13) and flat, so as to minimise the focal variation across the imaged regions. This can most easily be achieved by fixing cells as a monolayer. Cell deposition as monolayers provides additional advantages, including a maximal surface/volume ratio, thus facilitating shorter dehydration and fixation times. Further, we wanted to avoid the long incubation periods (overnight) commonly used to fix sperm droplets on poly-lysine coated coverslips in a wet chamber (14, 15). Such long incubations can result in artefacts from sample degradation (Fig. S1), and increases the processing time to over a day. Further such long incubation times can lead to salt precipitation from the buffer and surface contamination. To avoid this, cells are typically washed multiple times through centrifugation. These washing steps, and notably for sperm which are fragile, have a detrimental effect on the cellular integrity and induce visible mechanical damage (Fig. S1).

To circumvent these problems, we have developed a protocol where diluted – but not washed – sperm formed a robust monolayer on a gold-coated surface after a very short incubation (~1 min). The short incubation avoids the above mentioned exposure to shear forces, providing excellent preservation of cellular and non-cellular structures. Attachment is achieved via thiol groups of the cysteine residues in cell surface proteins and allows a covalent attachment to gold-coated surfaces (16). The strength of the attachment is sufficient to prevent the detachment of the cells throughout the fixation and multi-step dehydration. The thin gold layer can simply be formed through sputter deposition or thermal evaporation and is compatible with a variety of substrates (silicon wafers, coverslips, mica sheets, etc.) (13). Of additional advantage, the gold layer is conductive, meaning it is possible to minimise the thickness of the final conductive coating on cells required to prevent charging induced by the exposure to the electron beam. When the cells are in direct contact with a conductive surface, this coating layer (typically carbon or metal) can be thinner than in traditional SEM methods, because it needs only to prevent a local charging on the cell surface (Fig. 1C).

**Figure 1.**
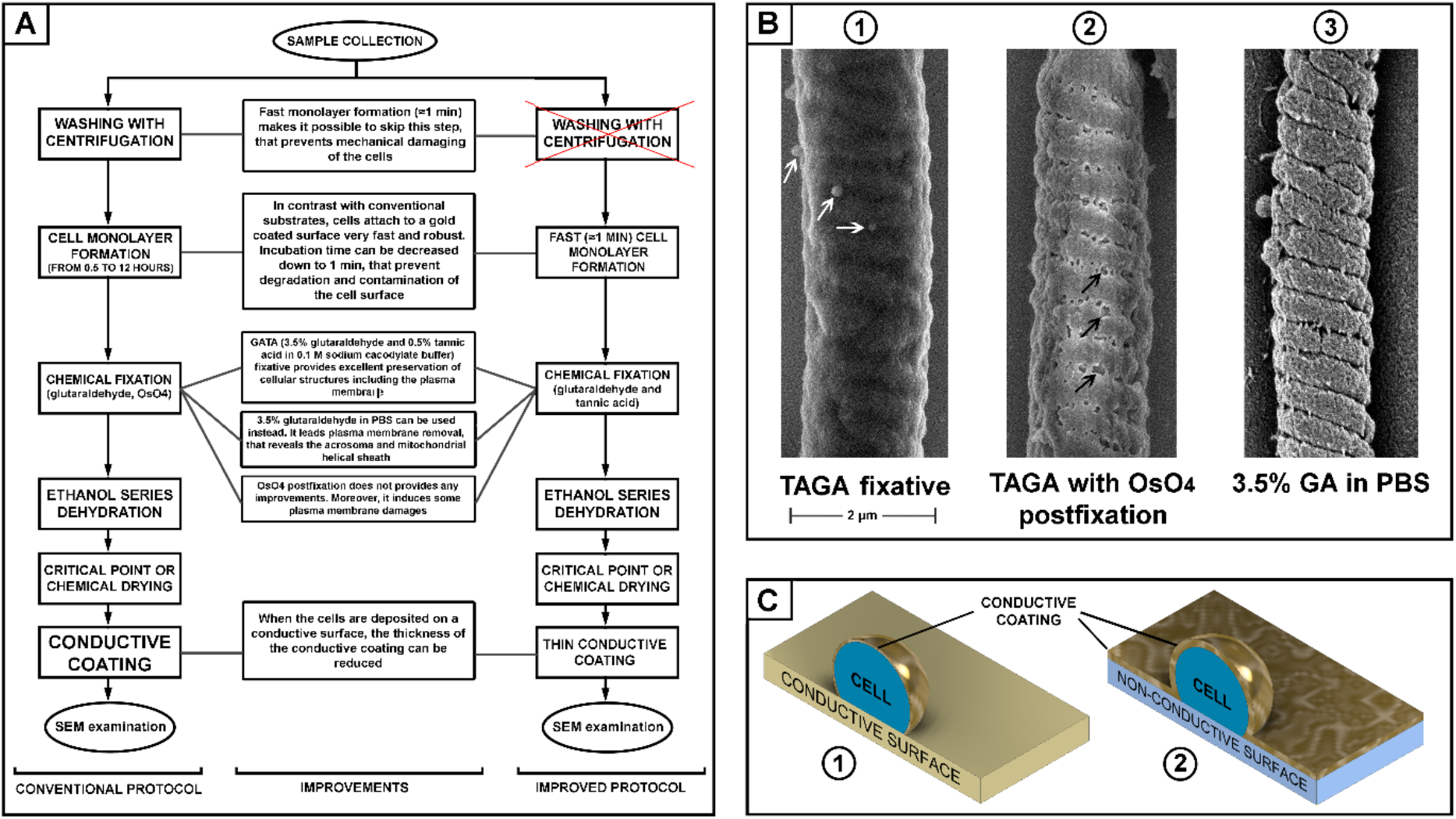
(A) Scheme of sperm preparation for SEM examination – conventional and improved protocols. (B) SEM pictures of a mouse sperm mid-piece with different fixation: glutaraldehyde and tannic acid (B1, microvesicles are indicated with white arrows); glutaraldehyde then post-fixation with osmium tetroxide (B2, plasma membrane damage is indicated with black arrows); and fixed with glutaraldehyde in PBS (B3, removal of the plasma membrane revealing the helically arranged mitochondria sheath), scale bar 2 μm. (C) A schematic illustration of the advantage of a conductive substrate (C1) for SEM sample preparation – the conductive coating of the cells can be thinner than it should be for a non-conductive substrate (C2).

We also aimed to test the relative merits of several commonly used fixatives. Buffered glutaraldehyde is the most common fixative in electron microscopy. It provides fast and reliable fixation of cells cross-linking their proteins by the dialdehyde (17). Osmium tetroxide and/or tannic acid are then commonly used during the second step of fixation (“post-fixation”) to cross-link and stabilise lipids in cellular membranes (18). Tannic acid can be used in mix with glutaraldehyde (TAGA fixative) to fix cellular proteins and lipids simultaneously (19). Comparing different fixative and combinations, we found that a mix 0.5% tannic acid and 3.5% glutaraldehyde in 0.1 M sodium cacodylate buffer provided the best preservation of the sperm plasma membrane. Using this fixative, we found that osmium tetroxide post-fixation, which is a common practice in many standard protocols (20, 21), actually induces damage to the plasma membrane. Thus, this post-fixation step should be avoided to prevent compromising the quality of the sample (Fig. 1).

The main purpose of fixation is to preserve the morphology of cellular structures as close as possible to their native conditions. Nevertheless, we identified research advantages from the removal of the plasma membrane. Specifically, we found that fixation using 3.5% glutaraldehyde in PBS without osmication resulted in the removal of the plasma membrane from the majority of sperm, while still providing good preservation of the internal organelles. In doing so, it is possible to reveal structural features of sperm that would be difficult to appreciate in standard SEM or TEM protocols. These included the helically arranged mitochondrial sheath and the relationship between the acrosome and other regions of the sperm head, which is not visible on SEM micrographs. Except for the use of 3.5% glutaraldehyde/PBS, the processing of samples was identical to that described above.

Most importantly, TAGA fixative provided a preparation compatible with high-quality SEM imaging of mouse sperm. Visible structures included the falciform head of the mouse sperm and a clear impression of each of the helically arranged mitochondria underneath the plasma membrane in the mid-piece, the fibrous sheath within the principal pieces, and the annulus between the two regions (Fig. 2). The plasma membrane was intact with no sign of degradation or surface damage. Additional sub-regions of the sperm head became clearly visible, including the equatorial region and the head-tail coupling apparatus (Fig. 2B). Interestingly, both of these structures had multiple vesicles associated with them (Fig. 2B-C, arrows). While the identity of these vesicles was not tested, we hypothesise that they are epididymis-derived exosomes, known as epididymosomes, in the process of fusing with the sperm. This process is known to be critical for sperm function, but it is still poorly understood (22, 23). The distinct alignment of these structures along the equatorial segment (which corresponds to the site of sperm-oocyte fusion (24)), and the head-tail-coupling apparatus (at the junction of the sperm head and tail), is supportive of the targeted transport of molecules from epididymal epithelial cells to the transcriptionally and translationally silent sperm to modify sperm function via a broad set of processes collectively known as epididymal maturation. These observations highlight an additional advantage of the current method, and specifically the avoidance of washing, in that such structures, have not, to the best of our knowledge, been observed previously.

**Figure 2.**
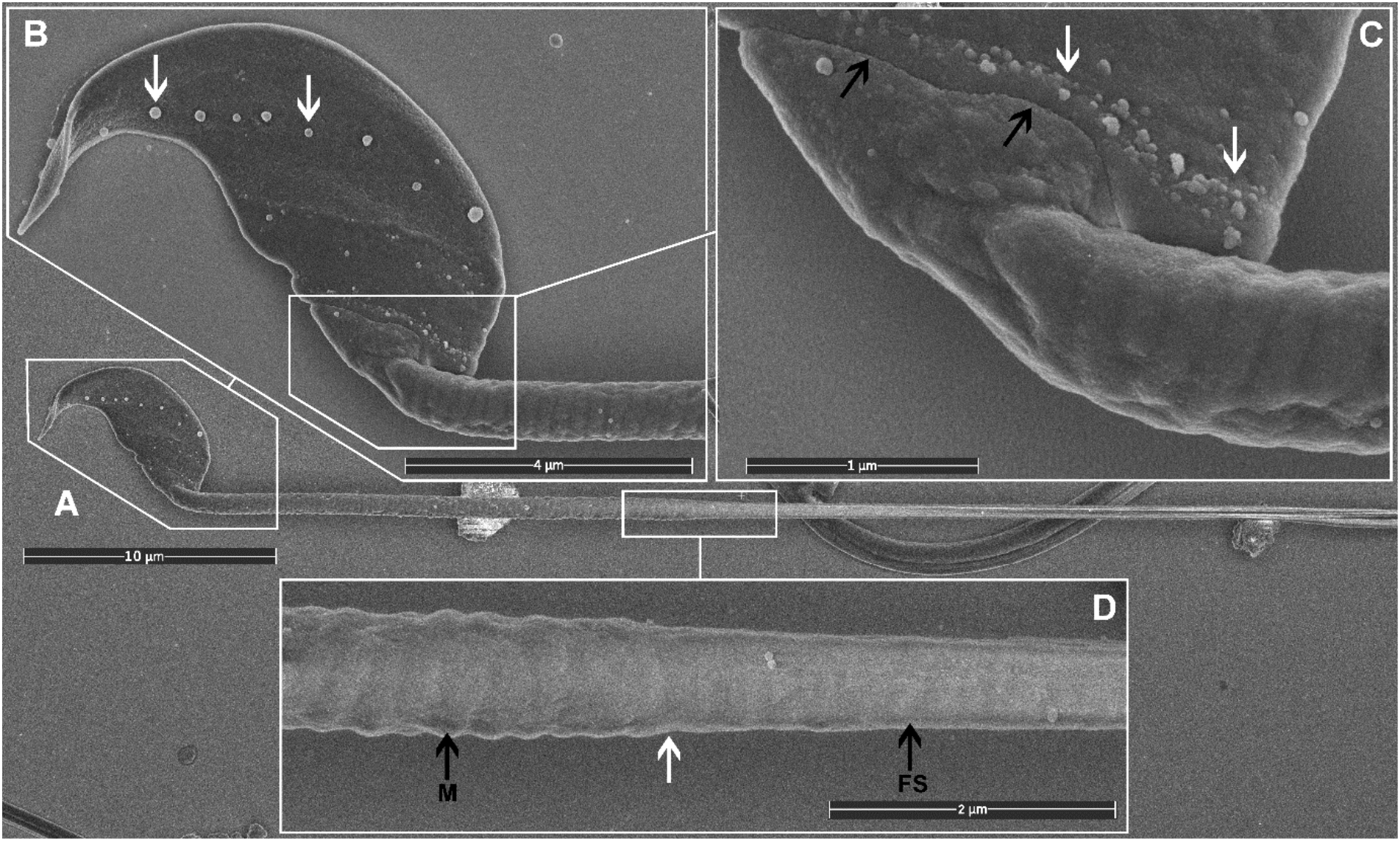
SEM images of mouse sperm fixed using TAGA. (A) A single sperm at low magnification. (B) The head structure. The arrows mark the presence of exosome-like structures decorating the equatorial region. (**C**) The sperm head-tail-coupling apparatus (black arrows) where the sperm head is attached to the flagellum. The white arrows mark the presence of a second line of exosome-like structures attached to the plasma membrane. (D) The sperm mid-piece principal piece junction (white arrow). A clear impression of the helically arranged mitochondria (M) is visible on the left, and the fibrous sheath (FS) of the principal piece on the right.

As indicated above, within the context of sperm biology, there are occasional advantages of removing the plasma membrane. Here, we show that this effect can, in fact, reveal previously unappreciated ultrastructural detail within sperm (Fig. 3). As shown in Fig. 3, the removal of the plasma membrane revealed the architecture of the helically arranged mitochondria in the mid-piece, the ring structure of the annulus at the junction of the mid- and principal pieces, and the whale boning-like structure of the perinuclear theca immediately below the acrosome within the sperm head. Further, within the principal piece you can clearly see the circumferentially arranged ribs of the fibrous sheath and for the first time, we believe, the visualisation of the fibrous longitudinal structures associated with the fibrous sheath (Fig. 3C). While the identity of these structures is currently unknown, they are consistent with the quadrilaterally-arranged domains where Catsper channels have been localised using super-resolution microscopy (25). Such structures would be virtually impossible to visualise using standard TEM methods and masked using standard SEM methods.

**Figure 3.**
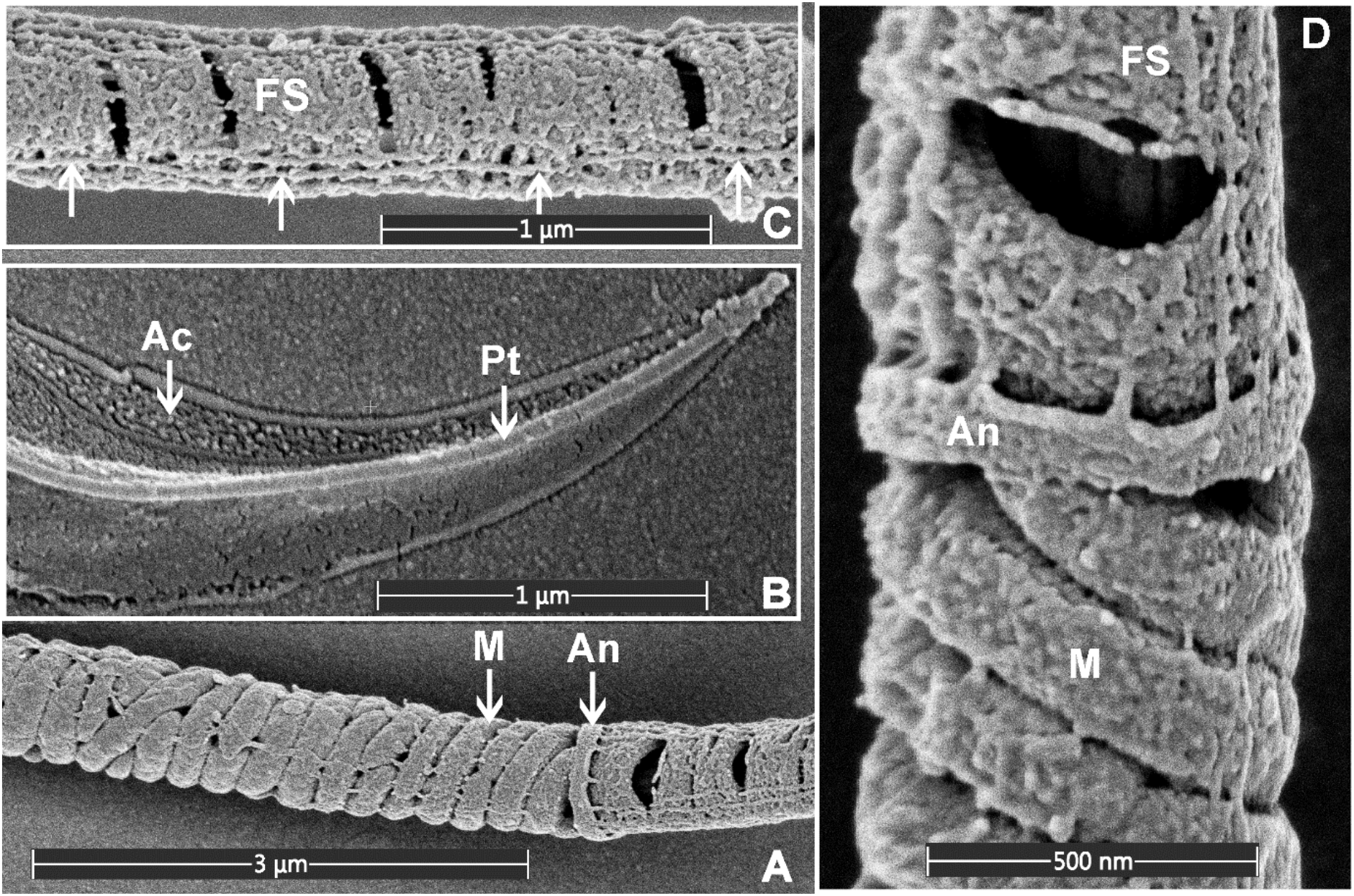
SEM images of mouse sperm fixed using 3.5% glutaraldehyde in PBS without post-fixation. (A) The helically arranged mitochondria (M) within the mid-piece and the annulus (An) which prescribed the junction between the mid- and principal pieces of the sperm tail. (B) The apical tip of the sperm head, revealing the acrosome (Ac) and perinuclear theca (Pt). (C) The principal piece of the sperm tail revealing the circumferentially arranged ribs of the fibrous sheath (FS) and longitudinal structures running along the length of the principal piece (marked with the arrows). (D) A higher power image of the junction between the mid- and principal piece junction of a sperm tail.

### 2. Internal sperm morphology

To examine the external morphology of a cell with SEM, the biological sample should be fixed, stained with heavy metals, and embedded into resin. Then thin layers of resin should be removed from the face of the block with an automated ultra-microtome, and the surface scanned with electrons (26, 27). Here we have modified our rapid sperm processing protocol to facilitate such imaging. Cryogenic preservation by high-pressure freezing (HPF) was chosen as the method as it has been repeatedly shown to result in the best morphological preservation of cells and tissues (28–30).

We have developed an original HPF-FS based protocol that optimises cell concentration and provides the ability to collect images of full sperm sections (and a large number of cells per sample) via SEM backscatter imaging. Sperm were processed as described in the Materials and Methods section. The resin block was pre-trimmed, coated with 10 nm of carbon, and imaged with backscattered electrons (Figures 5, 6). The data collection can be conducted from a large surface area, containing many full cell cross-sections with high-resolution, in an automated fashion, using dedicated software, such as MAPS (Thermo Scientific). Although serial block-face imaging can be performed if the three-dimensional architecture is required, we found that even a single cross-section can be extremely informative as it will allow an appreciation of the internal connections between, for example, the sperm head and tail (Fig. 6). Such images can be obtained on any conventional SEM and data collection is fast enough to provide, for example, statistically significant data on the morphology of unstudied sperm species or the consequences that specific genetic mutations have on sperm ultrastructure.

The practical resolution limit for examination of such samples with SEM in backscattering electrons mode is approximately 5 nm. In cases where a higher resolution is required, the embedded cells can be sliced for conventional TEM examination.

### 3. FIB-SEM tomography – full cell 3D reconstruction

Sperm, like other cells, are complex three-dimensional structures. As such, there will be numerous advantages of being able to image them in three-dimensions at electron microscopic resolution.

In comparison to conventional SEM and TEM visualisation, collection of FIB-SEM data is a slow process and can take hours to days per cell. This process consists of two parts: ion milling and imaging. While the imaging time is essentially defined (and limited) by the staining and the desired resolution, the milling time in FIB-SEM tomography strictly depends on how far the region of interest extends from the face surface of the block (distance from the top surface). Monolayer embedding, and thus a known cell orientation, makes it possible to minimise the milling depth (Fig. 5, 7), leading to a dramatic reduction in the milling time required. This is especially true for spermatozoa which typically are not more than 2 μm in thickness. Further, in the methods section, we describe a double osmication staining protocol which delivers an improved contrast of internal sperm structures, thus reducing the SEM visualisation dwell time. In addition, this protocol makes cells dark enough to be visible under a conventional light microscope, thus greatly facilitating the identification of target the area of interest for ion milling (Fig. S2). Finally, monolayer preparation also facilitates the pre-orientation of the samples to maximise the information contained in each slice regardless of the chosen imaging modality (TEM, SEM or FIB-SEM). FIB-SEM tomography provides a three-dimensional structure of the examined volume with nanometer-scale voxel size (Fig. 8A). Such graphical data can be analysed through segmentation and virtual cross-section to reveal external and internal morphological features (Fig. 8B-D). For example, it is possible to appreciate the articulation between the sperm head and tail at the HTCA, the arrangement of the acrosome relative to the nucleus, and even the presence of individual mitochondria, the outer dense fibres and fibrous sheath.

## Discussion

Sperm are complex cell machines that have been sculpted by evolution to optimise the chances of fertilisation. Their movement and hydrodynamics are critically dependent on the mechanics of the whip-like sperm tail and influenced by the shape of the sperm head. Within polyandrous species wherein female mate with multiple males, data suggests that small changes in these structures are the difference between fertilisation success or failure in sperm competition. As such, sperm structure is a major determinant of evolutionary change (1). Conversely, mouse sperm are composed of more than 2000 different proteins, defects in any of which could, in theory, lead to infertility (31).

Nevertheless, despite this, sperm are rarely visualised in three dimensions, and almost always electron microscopy of sperm is limited by a conventional SEM visualisation of the external morphology and a conventional TEM using protocols developed in the 1970s. Accordingly, there is a clear need to develop rapid and contemporary sperm processing techniques. To fill this void, we have developed three optimised protocols for SEM visualisation of sperm – from an improved conventional SEM to what we believe is the first example of FIB-SEM technique for sperm capable of full-volume reconstruction with nanoscale resolution. We show that a gold-coated surface is an ideal substrate for preparation of sperm monolayers for SEM examination. Covalent adhesion of sperm to such surfaces is very fast and robust and is compatible with a range of common surfaces including coverslips, slides, silicon wafers using a conventional sputter coater. Such rapid attachment avoids mechanical damage and allows the visualisation of previously unappreciated structures both on and within sperm. Herein we have used mouse sperm as a test case but predict the same protocols will be of broad utility across species.

In the sperm preparation for the external morphology examination, the fast-covalent attachment between the sperm cells and the gold-coated substrate also makes superfluous any washing step based on centrifugation, thus minimising mechanical damage and preserving fine structures, including microvesicles associated with sperm plasma membranes. Further, via a simple manipulation of a standard fixation protocol, using glutaraldehyde in the absence of osmium tetroxide post-fixation, we have shown that “naked” sperm can be produced and visualised (Fig. 1, 3). Through this protocol, we were able to visualise several previously unappreciated substructures, including fibrous longitudinal structures running along the length of the fibrous sheath. Further, the developed protocol for internal sperm morphology visualisation provides a unique possibility to collect full axial cross-section pictures for multiple cells in a relatively short time with a TEM-like quality (Fig. 6).

FIB-SEM tomography is amongst the most informative techniques in modern room temperature electron microscopy, but it is an extremely time-consuming method. The developed protocol provides a significant improvement in data collection speed through minimisation of the depth of ion milling and optimisation of targeting of the area of interest using light microscopy (Fig. S2). We believe this is the first example of three-dimensional ultrastructural analyses of the sperm. FIB-SEM allows the full-volume reconstruction of cells; thus, it can dramatically improve the field’s understanding of the structure-function relationship in sperm, including allowing accurate three-dimensional topology of relevance to hydrodynamics research and ultimately the understanding of sperm competition.

Collectively, these three rapid protocols have the potential to accelerate discovery in the relationship between sperm structure and function, including the analysis of the consequences of gene modification or environmental exposure. The choice of ultrastructural technique to be used will be dependent on the scientific question and available equipment and is summarised in Table 3.

## Materials and Methods

Within this section, we describe the methods and protocol developed to allow high-resolution visualisation of sperm internal and external morphology. While several methods were tested, only the optimal methods are described as a step-by-step instruction. These are complemented with commentary on where particular care should be taken to ensure the attainment of high-quality images.

### Reagents

- 1X Phosphate Buffered Saline (PBS)
- Glutaraldehyde (Sigma, G5882)
- Ethanol (Merck, 100983)
- Acetone
- Osmium tetroxide
- Tannic acid
- Epoxy embedding medium (Epon 812, NMA, DDSA, BDMA)
- Epoxy glue
- Embedding capsules (8 mm, flat-ended)
- Carbon Paint (Ted Pella, DAG-T-502)
- Mouse sperm

### Equipment

- Coverslips (can be replaced with silicon wafer or mica sheets)
- Micropipette capable of dispensing 30 μl
- Plastic transfer pipettes, 3 ml
- SEM Specimen Stubs (12.7 mm, Ted Pella, 16111)
- Manual glass cutter
- Sputter coater (e.g. BAL-TEC SCD 005 or similar)
- Critical point dryer
- High-pressure freezer
- 6 mm sapphire disks (0.1 mm thickness)
- 6 mm aluminium carriers for high-pressure freezing (0.3 mm/flat and 0.05/0.15 mm)
- Automatic freeze substitution unit
- Benchtop centrifuge capable of 100 × g with 1.5 ml centrifuge tubes
- Ultra-microtome with a diamond knife
- Thermal evaporating carbon coater
- Scanning electron microscope (e.g. FEI Helios G4 or similar)

### Image Processing

The FIB-SEM data stacks were processed and visualised using Dragonfly 2020.2 (Computer software). Object Research Systems (ORS) Inc., Montreal, Canada, 2020; software available at http://www.theobjects.com/dragonfly.

### Sperm sample collection

Animal procedures were performed in accordance with Australian National Health and Medical Research Council (NHMRC) Guidelines on Ethics in Animal Experimentation and approved by the Monash University School of Biological Sciences Animal Experimentation Ethics Committees. Sperm cells were collected from the cauda epididymides using the back-flushing method (33) then diluted into PBS to approximately 10^7^ ml^−1^. Care was taken to avoid shear stress from pipetting, which can result in sperm decapitation or tail damage.

### Sample preparation

#### 1. Samples for external morphology examination

*Substrate preparation* – coverslips were cut into 5×5 mm pieces with a glasscutter. The surface should be as clean as possible. Coverslips can be replaced with silicon wafer or mica sheet. Coverslip fragments were then coated with a gold layer using a sputter coater to a thickness of 30 to 50 nm. The thickness of the coating is determined by sputtering time and dependent on the sputter coater. Typical gold deposition rate for a conventional low-voltage sputter is from 5 to 10 nm/minute (34).

*Preparation of a sperm monolayer* - to prepare samples for external morphology examination, 0.1 ml of the sperm suspension in PBS was gently diluted in 1 ml of 0.1 M sodium cacodylate buffer, then a 30 μl droplet of this suspension was placed onto the fresh gold-coated substrate and sperm allowed to settle for 1 min, followed by washing with 0.1 M sodium cacodylate buffer for three times. The adhesion of sperm to the gold-coated substrate is mediated via the covalent attachment of cysteine-containing proteins to the gold-coated surface (16). For optimal adhesion, the gold layer should be fresh i.e. less than 24 hours from the coating.

*Chemical fixation* – a 30 μl droplet of fresh 3.5% glutaraldehyde and 0.5% tannic acid solution (TAGA fixative) in 0.1 M sodium cacodylate buffer was placed onto the cell monolayer immediately after washing to avoid drying. Cells were fixed for 15 min; then the fixative was removed via washing with 0.1 M sodium cacodylate buffer. To prepare sperm monolayer with removed plasma membranes, TAGA fixative should be replaced with 3.5% glutaraldehyde in PBS (Fig. 1).

*Dehydration of the cell monolayer* by immersing in graded ethanol 30% 50%, 70%, 90%, 96%, 100% and 100% for 5 minutes each. Residual ethanol was removed with a critical point dryer. To visualise the cells with SEM, the sample was mounted on a standard metal SEM stub with conductive carbon paint, and then coated with a ~5 nm thick gold or carbon layer.

#### 2. Samples for internal morphology examination

A 30 μl droplet of sperm suspension (approximately 10^7^ ml^−1^) in PBS was placed onto the fresh gold-coated (~10 nm) sapphire disk and sperm allowed to settle for 10 min. The droplet was then removed with a pipette and the disc placed on the flat side of 6 mm aluminium carrier (0.3 mm / flat) and covered with the second aluminium carrier (0.05/0.15 mm, side 0.05 mm, soaked in hexadecene). Then this “sandwich” (Fig. 4) was mounted in the standard 6 mm sample holder and frozen with a high-pressure freezer.

**Figure 4.**
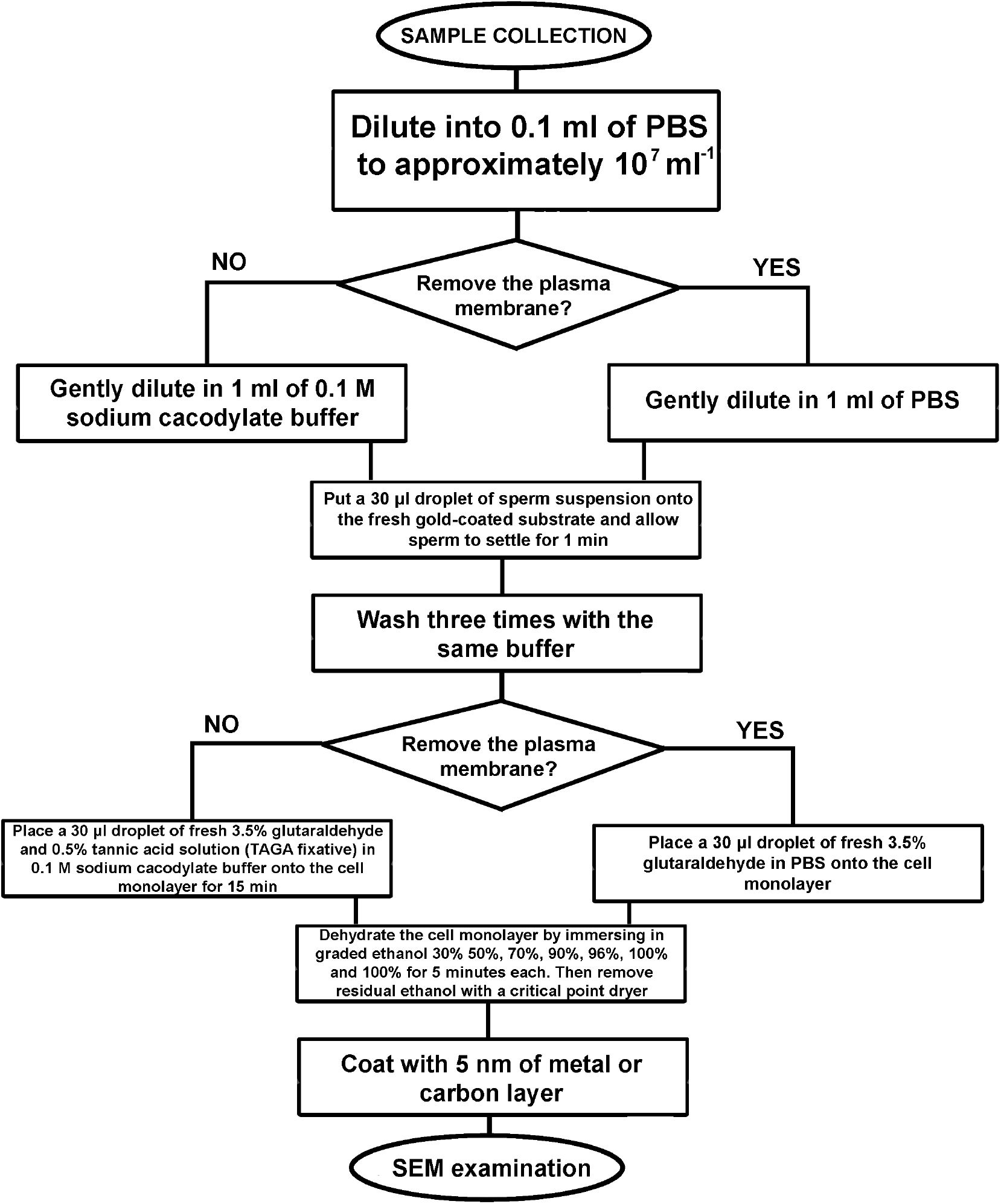
An algorithm of the developed sample preparation protocol for SEM examination of sperm external morphology.

Frozen samples were stored in liquid nitrogen and then put to individual cryogenic vials (1.5 ml) for freeze substitution using Leica AFS unit (Tab. 1).

**Table 1.** Comparison of the developed protocols.

| Method | Structural information, which can be collected | Time | Laboriousness and costs |
| --- | --- | --- | --- |
| Conventional SEM visualisation | External morphology with nanometre resolution, many cells can be examined, including automatic mode | Fast preparation and examination | Easy and cheap |
| SEM examination of carbon coated trimmed monolayer | Internal morphology (full cell cross-sections) with nanometre resolution, many cells can be examined, including automatic mode | Slow preparation and fast visualisation | Sample preparation is expensive and time-consuming. Visualisation is easy, fast and can be automatised. |
| FIB-SEM tomography | True three-dimensional data with nanometre resolution. External and internal morphological features simultaneously. | Slow preparation and very slow data collection | Very expensive and extremely time-consuming |

After freeze substitution and resin infiltration, the discs were put into 8 mm flat-ended embedding capsules for polymerisation using custom 3D-printed holders (Fig. 5). The holders keep the discs in the proper orientation during polymerisation.

**Figure 5.**
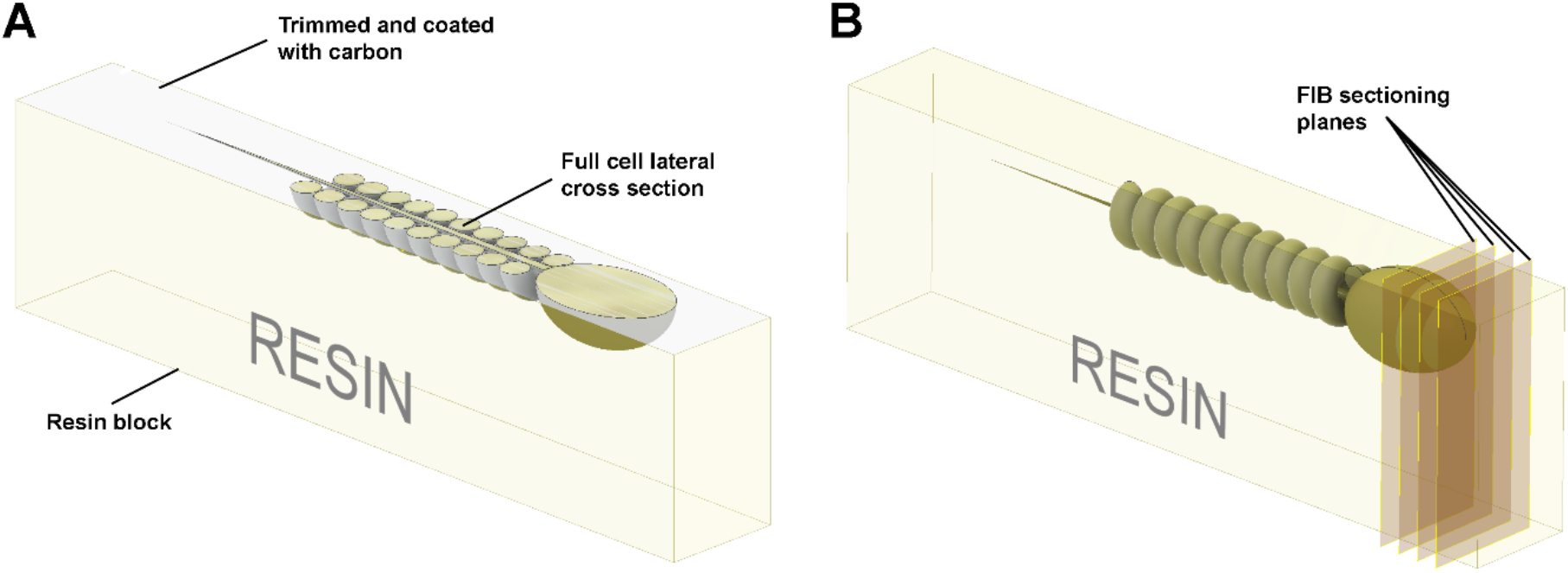
Schematic illustration of the preparation of resin embedded cell monolayer. The top surface is formed by detaching the sapphire disk from the resin block; thus, it is smooth, and the cells are localised precisely on the surface. (A) For the visualisation of full cell cross-sections, a layer of resin about half the thickness of the cell should be trimmed off with an ultra-microtome using a diamond knife, and the surface coated with ~10 nm of carbon. The cross-sections can then be visualised with SEM in backscattering electron mode. (B) To collect FIB-SEM volume data the ion beam serially removes thin layers from the block face surface.

To minimise the detachment of cells from the disc, the capsules were centrifuged at 100×g for 5 min before polymerisation. After polymerisation, the resin blocks were removed from the plastic capsules, then cooled with liquid nitrogen vapour to detach the sapphire discs (35). To prevent cracks forming during this operation, the resin should be fully polymerised via a curing time of 72 to 96 hours at 65 °C.

After detachment of the discs, the 6 mm resin blocks were each divided into nine parts with a fine blade (Fig. 6). Then each small block was individually glued to a SEM stub with epoxy adhesive (face side down, protected with a layer of parafilm), the top surface was polished parallel to the stub face, then the block was detached, flipped with the face side up and glued back (Fig. 7). The geometry of the block has a crucial meaning for precise trimming and block-face imaging; thus, the top surface should be made parallel to the stub.

**Figure 6.**
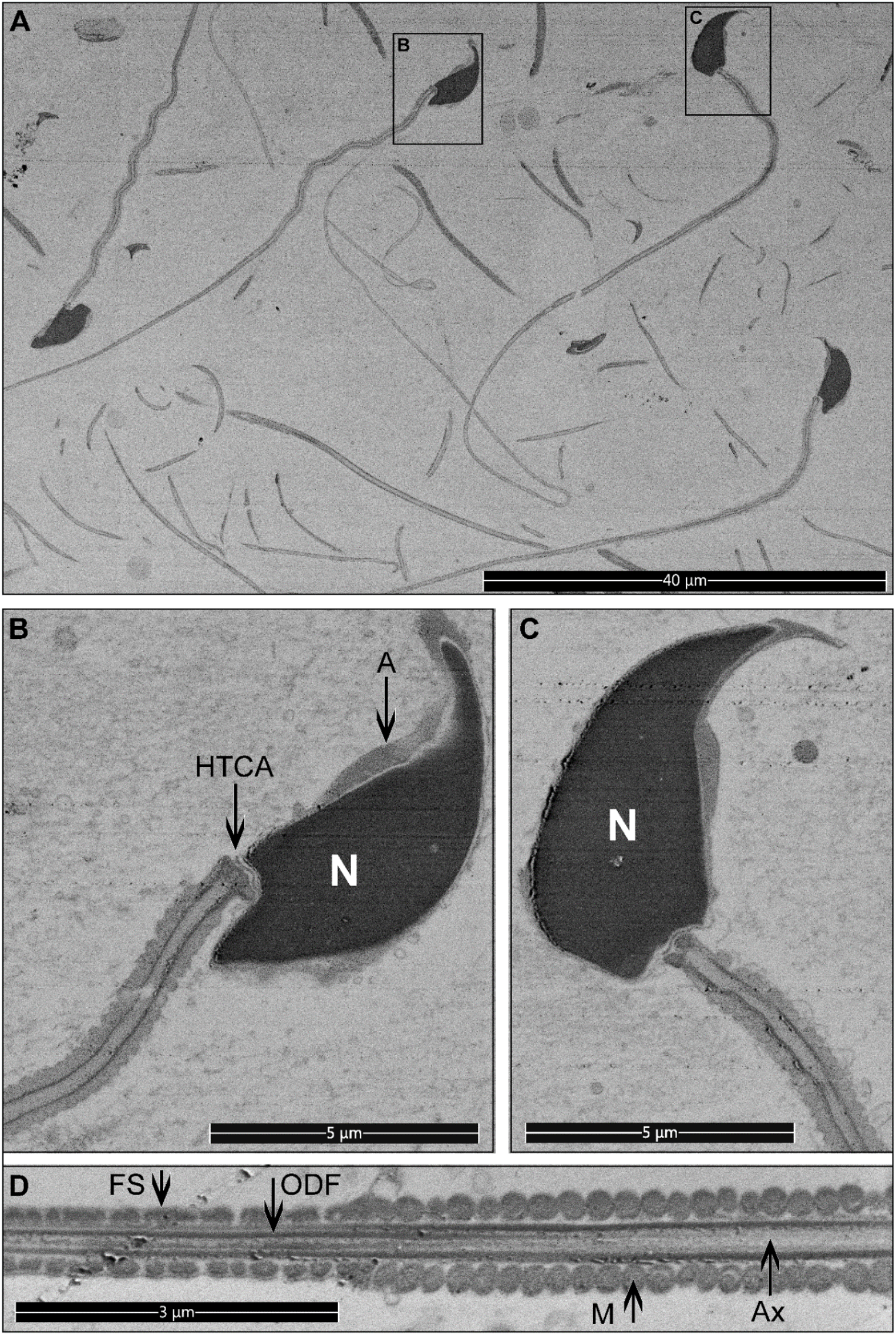
Longitudinal sections for mouse sperm. Pictures were collected from the top surface of a pre-trimmed block in backscatter mode. (A) Low magnification, multiple cross-sections of different sperm are visible. Scale bar 40 μm. (B-D) The internal structure of sperm cells. Scale bar 5 μm. M = mitochondria in the mid-piece. FS = fibrous sheath in the principal piece. ODF = outer dense fibers. Ax = microtubules of the axoneme. A = acrosome. N = highly condensed DNA within the nucleus. HTCA = head-tail-coupling apparatus.

**Figure 7.**
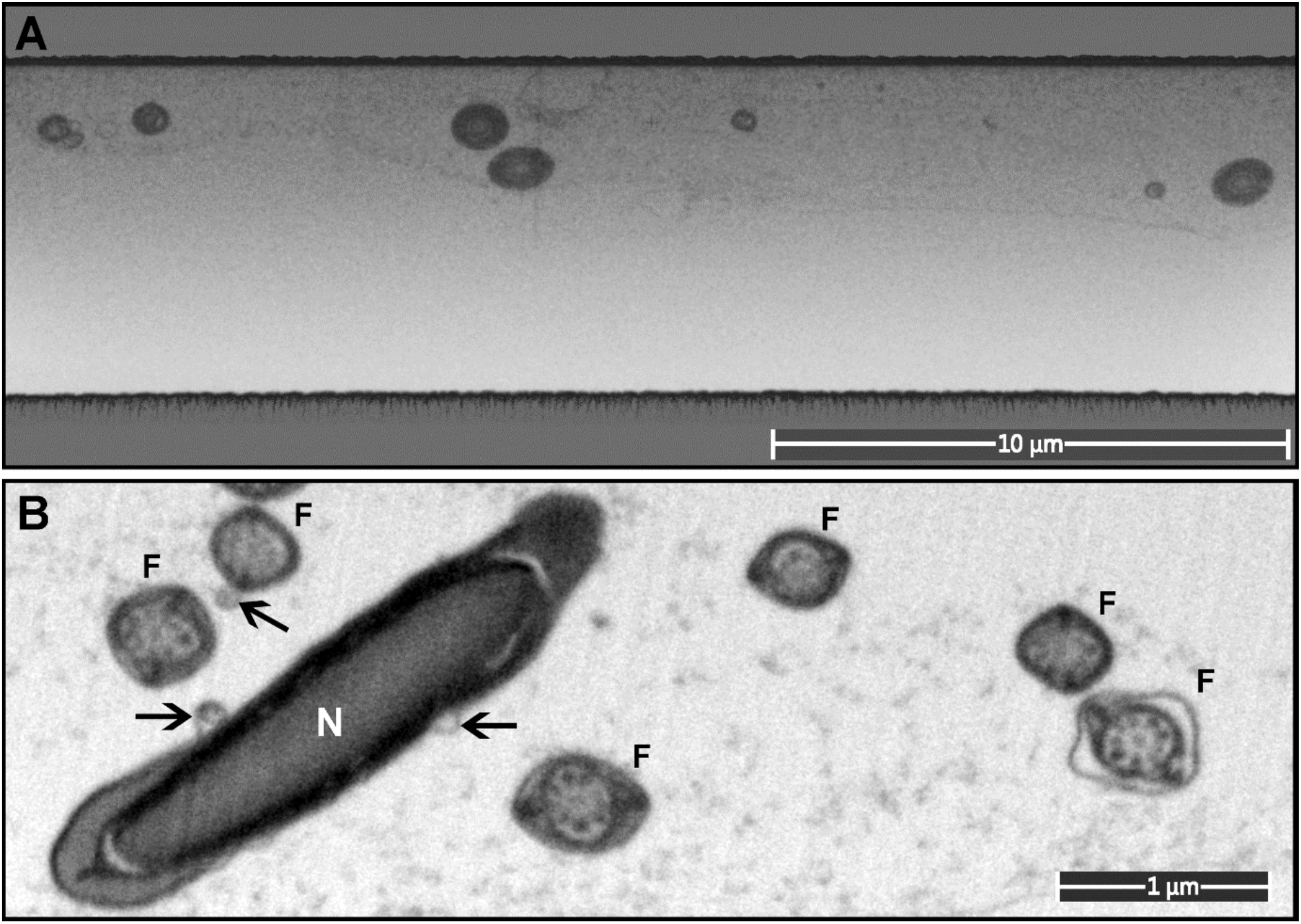
A monolayer of mouse sperm on the top surface of a resin block used for FIB-SEM. (A) An image of a trench on the surface of the FIB-SEM block (low magnification). (B) Cross-sections of sperm flagella (F) and nucleus (N). Exosome-like structures are marked with arrows.

A 0.5 μm layer of resin was precisely removed from the top surface of each sub-block using an ultra-microtome (step 50 nm) with a diamond knife to reveal the internal structures of the cells. Then the vertical sides of the block were coated with conductive carbon paint, and the top surface was coated with ~10 nm of carbon, and then examined with an SEM in backscattering electron mode.

High mechanical stability of the resin block surface is critical for precise sub-micrometer trimming. It was reached using hard Epon resin composition (Tab. 2).

**Table 2.**
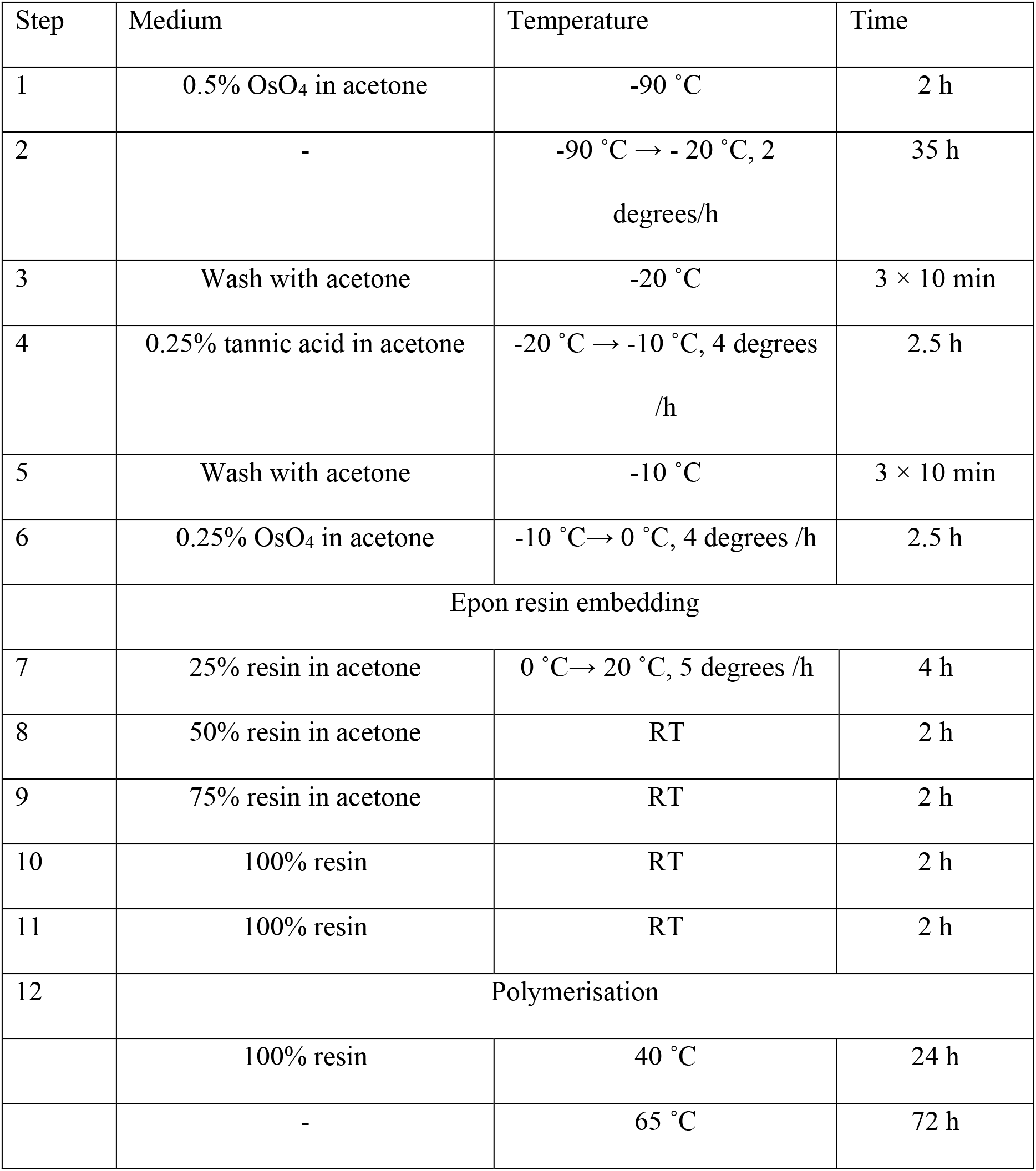
The protocol of freeze-substitution and resin embedding. RT = room temperature.

| Step | Medium | Temperature | Time |
| --- | --- | --- | --- |
| 1 | 0.5% OsO<sub>4</sub> in acetone | -90 °C | 2 h |
| 2 | - | -90 °C → -20 °C, 2<br>degrees/h | 35 h |
| 3 | Wash with acetone | -20 °C | 3 × 10 min |
| 4 | 0.25% tannic acid in acetone | -20 °C → -10 °C, 4 degrees<br>/h | 2.5 h |
| 5 | Wash with acetone | -10 °C | 3 × 10 min |
| 6 | 0.25% OsO<sub>4</sub> in acetone | -10 °C → 0 °C, 4 degrees /h | 2.5 h |
|  | Epon resin embedding |  |  |
| 7 | 25% resin in acetone | 0 °C → 20 °C, 5 degrees /h | 4 h |
| 8 | 50% resin in acetone | RT | 2 h |
| 9 | 75% resin in acetone | RT | 2 h |
| 10 | 100% resin | RT | 2 h |
| 11 | 100% resin | RT | 2 h |
| 12 | Polymerisation |  |  |
|  | 100% resin | 40 °C | 24 h |
|  | - | 65 °C | 72 h |

**Table 3.**
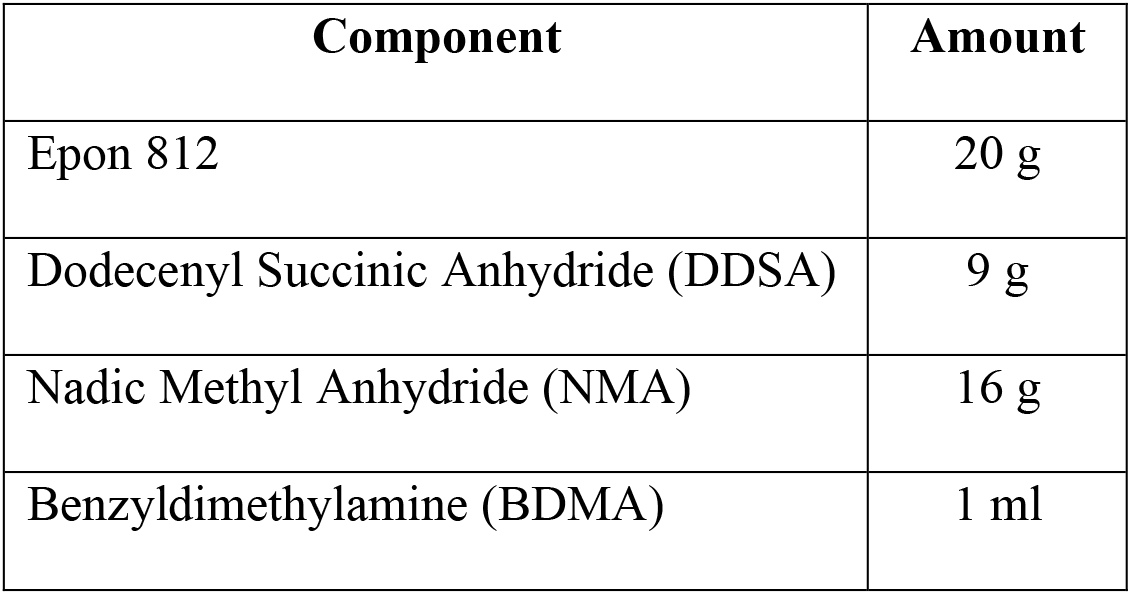
Composition of hard Epon resin.

#### 3. Samples for FIB-SEM tomography

The sperm samples for FIB-SEM tomography were frozen, substituted and embedded into resin as it was described above (Table 2). The blocks were glued to standard SEM stubs and prepared as described above (Fig. 2, 3) except trimming and carbon coating. Before FIB-SEM examination the blocks were coated with about 20 nm of gold.

### SEM examination

#### 1. External sperm morphology

The dry and conductive samples should be examined under high vacuum within the SEM to achieve the best resolution. To study the external morphology of the cells, the SEM was used in secondary electron detection mode. A FEG-SEM ThermoFisher Elstar G4 was used in this study, and the best quality of imaging was obtained with accelerating voltage of 2 kV, secondary electron mode (SE), and a work distance of 4 mm, operating in immersion mode with the through lens detector (TLD). The best signal/resolution was obtained when using a monochromated 25 pA beam probe and the dwell time 4 μs / pixel.

#### 2. Internal sperm morphology

To study the internal morphology of spermatozoa, the SEM was used in backscattering electron detection mode. For the FE-SEM ThermoFisher Elstar G4 used in this study, the best quality of imaging was obtained with accelerating voltage of 2 kV, backscattering electron mode (BSE), and a work distance of 3 mm, operating in immersion mode with the through lens detector (TLD). The best signal/resolution was obtained when using a 0.4 nA beam probe and the dwell time 8 μs / pixel.

#### 3. FIB-SEM tomography – full cell 3D reconstruction

The cells under the top surface of the block were localised using imaging with an electron beam at accelerating voltage of 30 kV. Then a layer of 500 nm platinum was precisely deposited above the targeted cells using a gas injection system (GIS). A trench (depth about 5 μm, **Fig.**8) was milled with gallium ions (30 kV, ion current 2.2 nA) on the border of the platinum spot; then the FIB-SEM data were collected using Auto Slice & View software (ThermoFisher). For the FE-SEM ThermoFisher Elstar G4 used in this study, the best quality of FIB-SEM imaging was obtained with accelerating voltage of 2 kV, backscattering electron mode, and a work distance of 4 mm, operating in immersion mode with the through lens detector. The best signal/resolution was obtained when using a 0.2 nA beam probe and the dwell time 6 μs / pixel.

**Figure 8.**
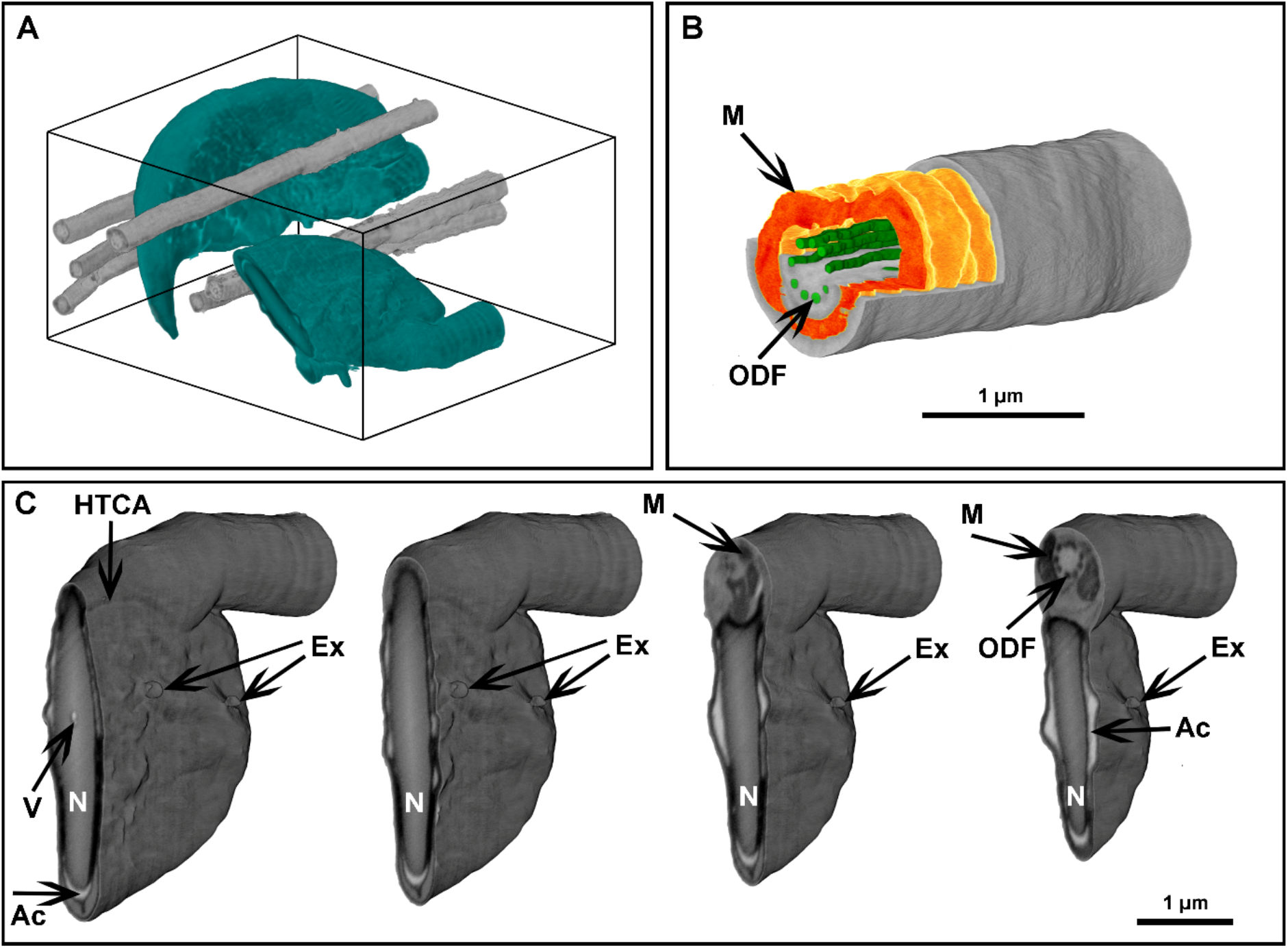
FIB-SEM tomography of a monolayer of mouse sperm. (A) Volume reconstruction 4.25 × 3.3 × 4.9 μm. Sperm head (green) and flagella (grey). (B) Segmented fragment of the mid-piece. (C) Virtual cross-sections (step 500 nm) of the head area. Ex = exosome-like structures attached to the plasma membrane. HTCA = head-tail-coupling apparatus. Ac = acrosome. M = mitochondria. N = highly condensed DNA within the nucleus. ODF = outer dense fibers. V = a potential nuclear vacuole. The figures are presented in orthographic projection.

**Figure 9.**
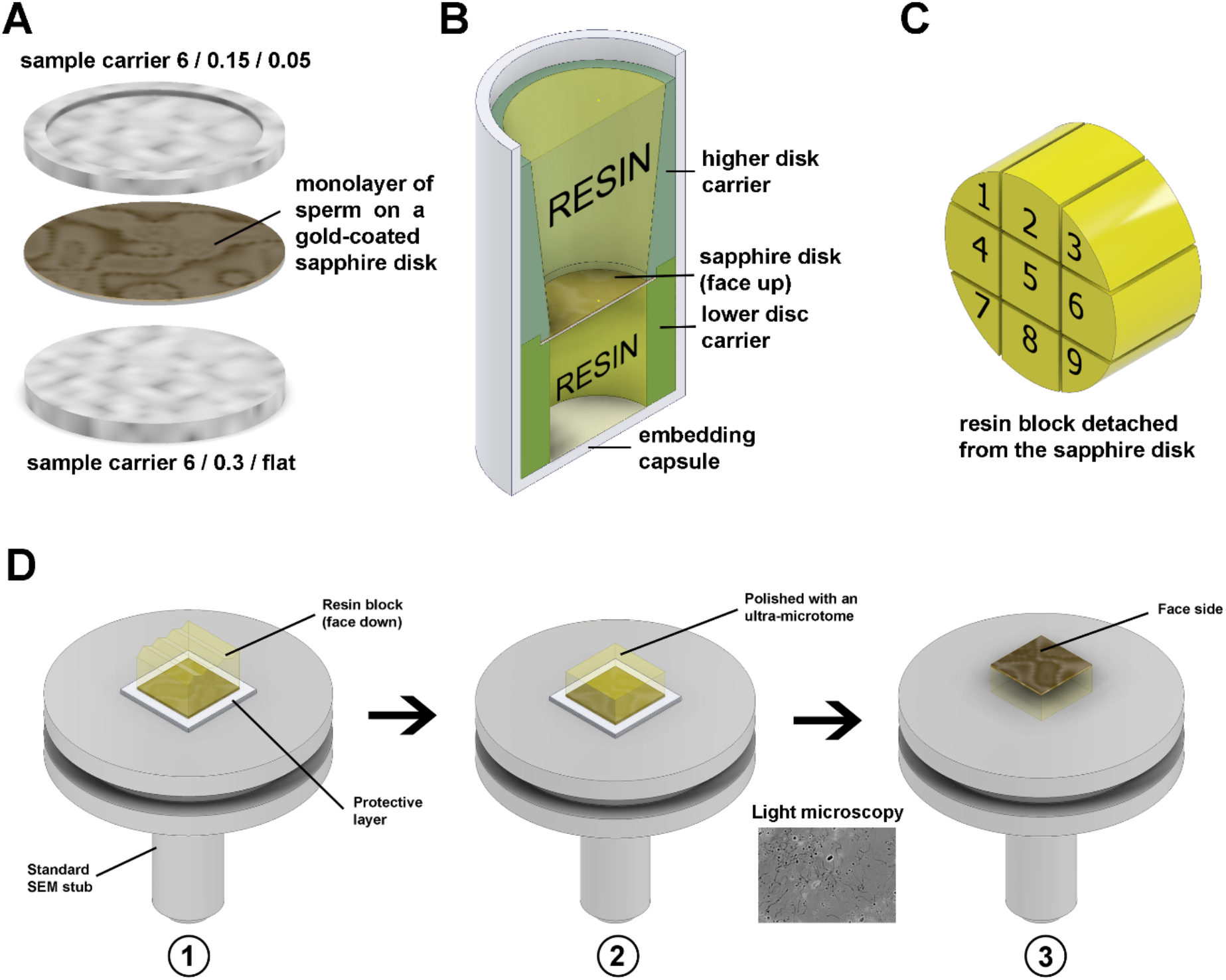
Freezing and resin block preparation. (A) Gold-coated sapphire disc with a monolayer of sperm between 6 mm aluminium carriers for high-pressure freezing. (B) Resin embedding into a plastic capsule using custom 3D-printed carriers. (C) Division of the 6 mm block to nine sub-blocks. (D) To make the top surface of the block parallel to the stub face it was glued to the stub with face side down, using parafilm as a protector (D1), then the top surface was polished parallel to the stub (D2), and then the block was detached, flipped and glued back with the face side up (D3). The thin gold layer on the face surface of the block is semi-transparent, thus after detachment from the stub, it can be examined under a conventional light microscope.

## Acknowledgements

We thank Dr Avinash Gaikwad for help with mouse sperm collection. We also thank the Monash Ramaciotti Centre for Cryo-Electron Microscopy.

## Authors’ roles

DK designed and conducted the experiments, wrote the article. DJM and GG conducted the experiments. AdM and MKOB supervised the project and edited the article and were involved in data analysis. The article was edited and approved by all authors.

---

**Figure S1.**
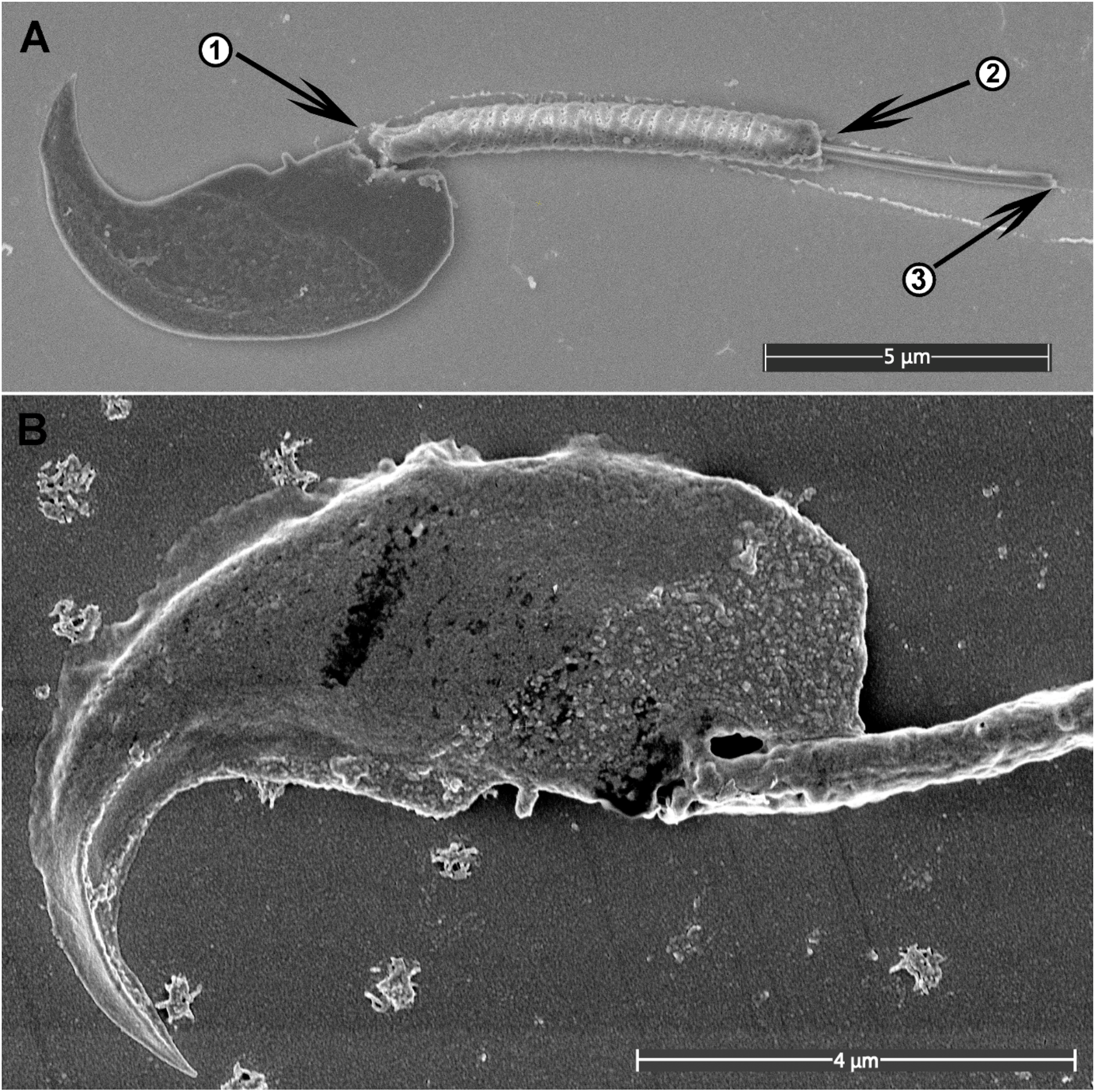
(A) Typical mechanical damages of sperm – rapture in the head-tail-coupling apparatus (1), snapping of the mitochondria sheath (2) and the tail (3). (B) Degradation of the cell surface as a consequence of prolonged incubation in glutaraldehyde (4 h, room temperature).

**Figure S2.**
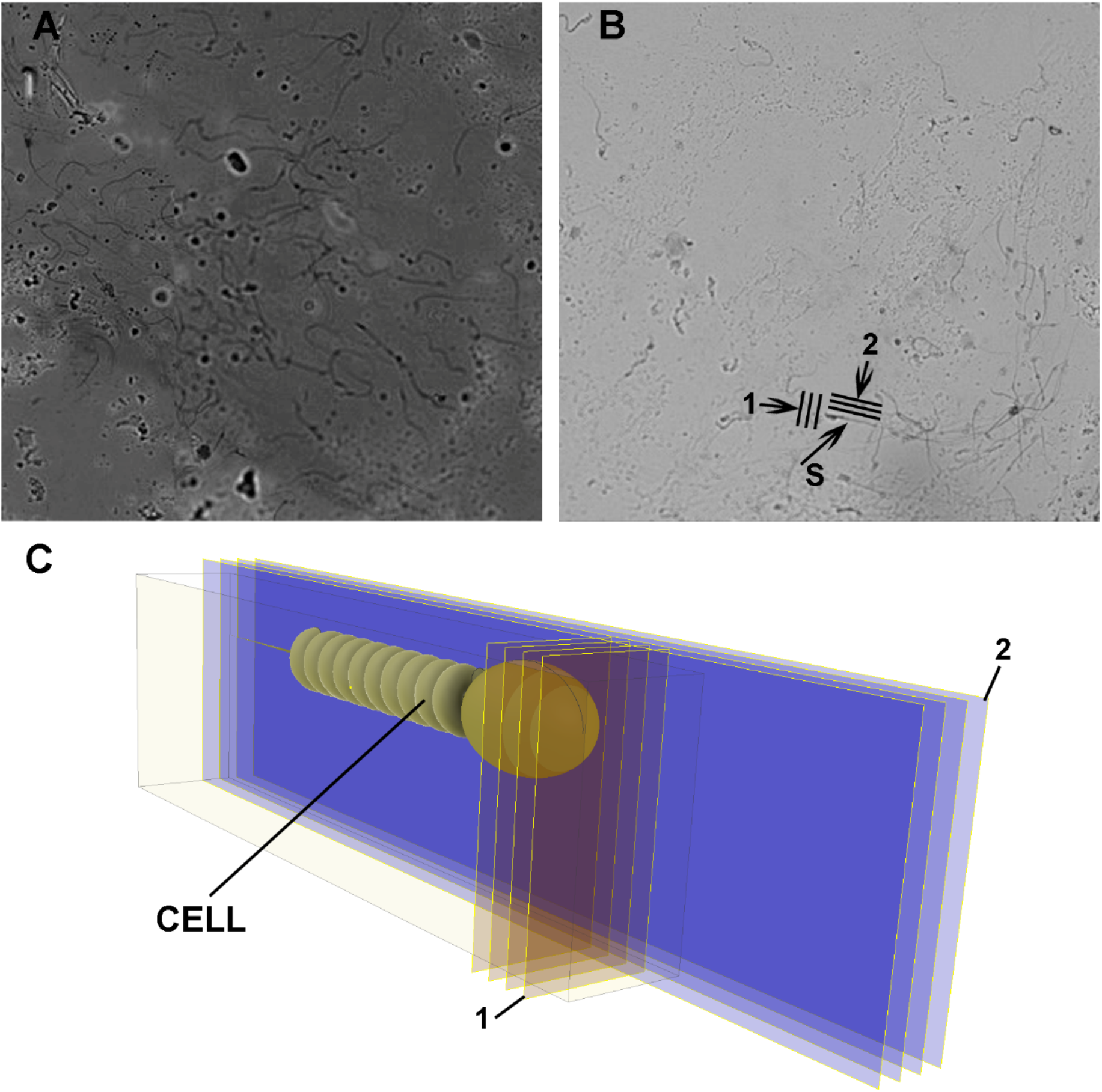
Cell targeting and orientation of FIB milling planes. (A, B) Light microscopy visualisation of the top surface of the resin block with embedded monolayer of sperm through a layer of gold (A) and from the area where gold is removed (B). The speed/quality rate is depending on the milling planes orientation. To reach the best resolution the milling plane should be orthogonal to the cell axis (B, C, 1). To achieve the fastest data collection with reduced resolution the milling plate should be parallel to the cell axis (B, C, 2). S = sperm.

**Figure S1.**
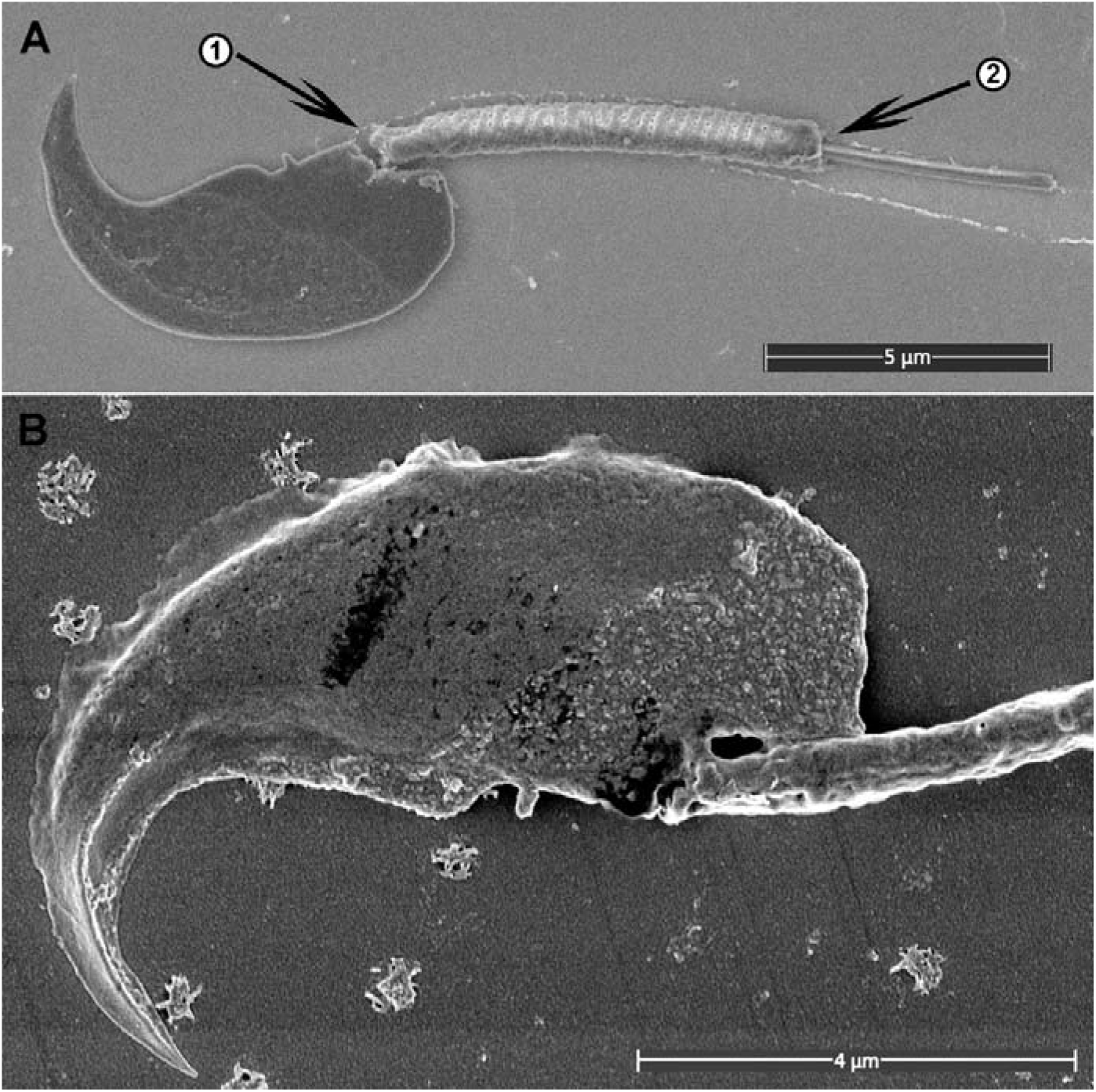
(A) Typical mechanical damages of sperm – rapture in the head-tail-coupling apparatus (1) and cell disaggregation (2). (B) Degradation of the cell surface with too long (4 h, room temperature) incubation in glutaraldehyde fixative.

**Figure S2.**
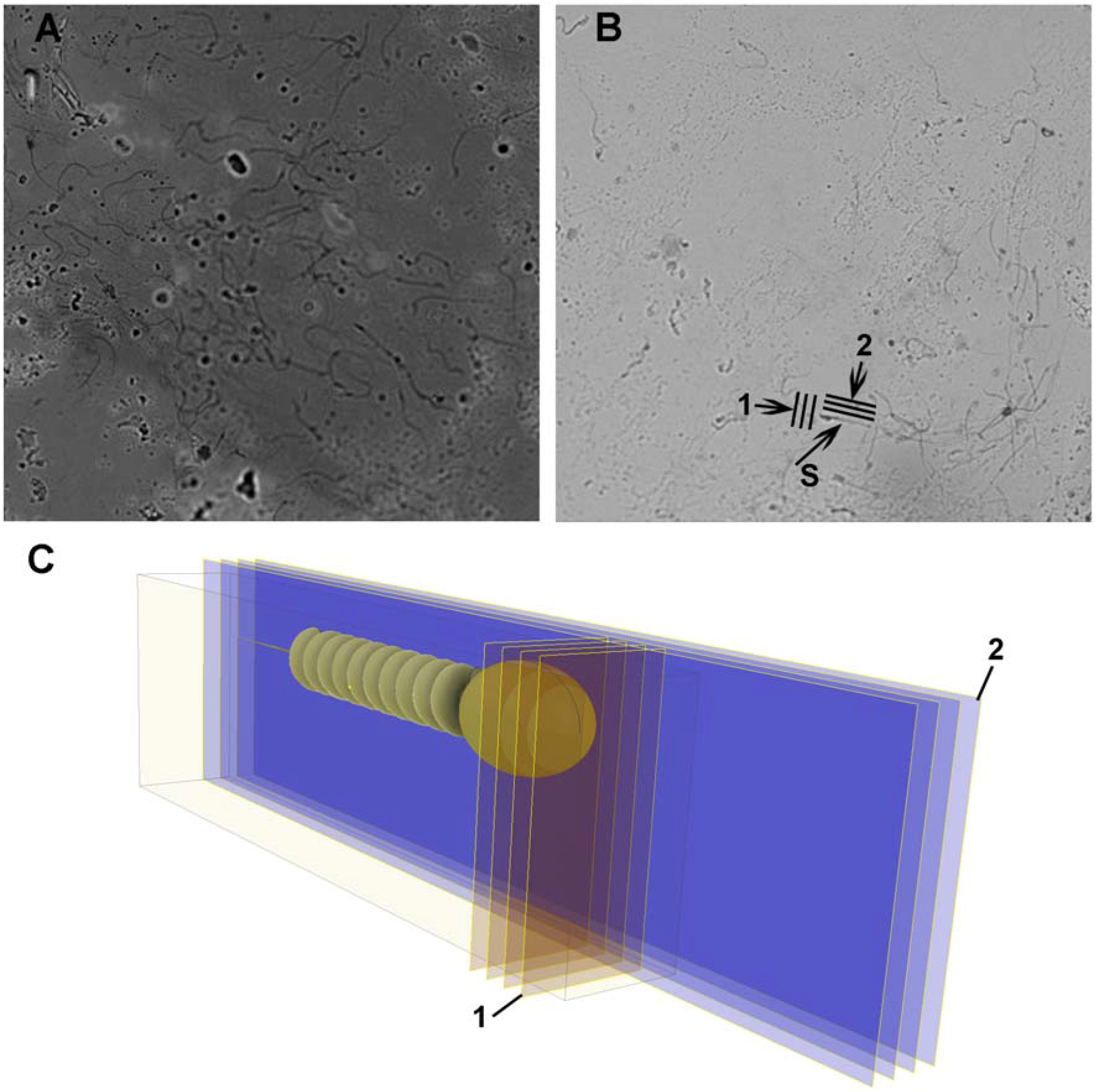
Cell targeting and orientation of FIB milling planes. (A, B) Light microscopy visualisation of the top surface of the resin block with embedded monolayer of sperm through a layer of gold (A) and from the area where gold is removed (B). The speed/quality rate is depending on the milling planes orientation. To reach the best resolution the milling plane should be orthogonal to the cell axis (B, C, 1). To achieve the fastest data collection with reduced resolution the milling plate should be parallel to the cell axis (B, C, 2). S = sperm.

